# Cargo levels control endosomal PI(3)P homeostasis through direct feedback between SNX17 and MTM1

**DOI:** 10.64898/2026.11.02.763636

**Authors:** Lewis Hancock, Ben Allouis, Anwar Sadat, Ruby Peters, Rebeka Butkovič, Alexander P. Walker, Francesca Bottanelli, Peter J. Cullen, Hannes Maib

**Affiliations:** University of Sheffield, School of Biosciences, Sheffield, UK; School of Biochemistry, Biomedical Sciences Building, University of Bristol, Bristol, UK; Institut für Biochemie, Freie Universität Berlin, Berlin, Germany

## Abstract

The sorting of internalised cell surface proteins through the endosomal network is crucial for cellular function. Phosphatidylinositol-3-phosphate (PI(3)P) is a critical lipid identity cue for organising endosomal sorting but there is limited understanding of how PI(3)P homeostasis is regulated. Here, we show that free PI(3)P is segregated into distinct endosomal subdomains and that depletion of the disease-relevant lipid phosphatase Myotubularin1 (MTM1) leads to its accumulation and mis-segregation, impairing cargo recycling. Complementary imaging and reconstitution approaches show that MTM1 recruitment and activity are controlled by binding to the endosomal cargo adaptor SNX17. This interaction is mediated by sorting motifs in the C-Terminal tail of MTM1 and is in competition with the same sorting motifs in cargo that bind to SNX17. This competitive binding mechanistically links cargo sorting to PI(3)P depletion and because SNX17 is itself a PI(3)P effector, establishes negative feedback regulating local PI(3)P levels. Importantly, we demonstrate that this feedback couples endosomal cargo levels to PI(3)P depletion. We propose that cargo levels thereby modulate endosomal phosphoinositide identity through cargo sensitive feedback between SNX17 and MTM1.

## Introduction

Membrane identity is crucial for cellular function, especially for the sorting of internalised transmembrane proteins (henceforth referred to as cargo) through the endo-lysosomal system. A crucial hallmark of this identity are the phosphoinositides, a small family of phospholipids generated by phosphorylation of their D-*myo*-inositol headgroup [1]. Following internalisation, receptors localise to sorting endosomes bearing the phosphoinositide PI(3)P [2]. This phospholipid is crucial for the recruitment of endosomal effector proteins (such as EEA1, Hrs and sorting nexins) that regulate cargo trafficking through binding to PI(3)P [3]. While EEA1 mediates cargo entry through fusion of early endosomes, Hrs mediates cargo degradation through the ESCRT machinery that assemble to form the degradative subdomain [4, 5]. In contrast, sorting nexins engage protein complexes such as Retromer, Retriever and Commander, which form the retrieval subdomain and concentrate cargo for recycling back to the plasma membrane or for retrograde transport to the Golgi apparatus [6, 7]. PI(3)P therefore sets the stage for endosomal sorting by establishing the membrane identity for both cargo entry and exit through retrieval or degradative domains.

The homeostasis of endosomal PI(3)P is regulated by two key players: the lipid kinase VPS34 complex II (VPS34/VPS15/Beclin 1/UVRAG) that generates endosomal PI(3)P, and the Myotubularin 1 (MTM1) lipid phosphatase that catalyses the reverse reaction and dephosphorylates PI(3)P back to PI [3]. When PI(3)P generation is blocked, endosome maturation is impaired due to a failure in the establishment of endosomal identity [8–11]. However, an excessive production of PI(3)P also leads to a failure in cargo recycling, with receptors failing to undergo endosomal exit [12, 13]. This is the case in X-linked centronuclear myopathy (XLCNM), a severe congenital myopathy caused by mutations in MTM1 and characterised by the endosomal accumulation of β1-integrin [14–16]. Additionally, MTM1 localisation to endosomes is sensitive to the metabolic state of the cell and during starvation regulates mTOR signalling and rearrangement of the endoplasmic reticulum [17, 18]. However, the molecular basis of how endosomal PI(3)P homeostasis is regulated to balance its generation to its depletion is still unclear. While it has been well established that Vps34 is recruited to endosomes via binding to active Rab5, [19–21] MTM1 recruitment and activity is less well understood.

The regulation of phosphoinositide conversions often includes complex product and substrate induced positive, negative or mixed feedback loops: these have been intensely studied for the dynamics of plasma membrane phosphoinositides [22–27]. Endosomal phosphoinositide conversion pathways are experimentally more difficult to assess but likely involve similar feedback mechanisms. A necessity of these feedback loops is that enzymatic activity is coupled to the functional outcome of a cellular process. For trafficking through endosomes this outcome is primarily the sorting of selected cargo proteins.

Far from being passive bystanders, cargo proteins are active modulators of their endosomal sorting [28, 29]. At the level of recycling, this is achieved via the presence of sorting motifs, such as the NPxY motif in the cytoplasmic tail of β1-integrin. This specific motif binds to the PI(3)P effector SNX17 for entry into the retrieval subdomain via the Retriever complex [30, 31]. Similarly, the PDZ binding motif in the glucose transporter GLUT1 binds to SNX27 for Retromer mediated entry into this subdomain [32]. On the flipside, ubiquitination of key lysine residues in EGFR leads to the recruitment of the ESCRT machinery and formation of multivesicular bodies and cargo degradation [33]. Cargo, therefore, plays an active role in the recruitment of key effector proteins and the amount of cargo present on an endosome will critically shape its functional state. However, until now there was no mechanistic detail of how cargo modulate the phosphoinositide identity of endosomes.

Here, we identify a feedback mechanism that links cargo to endosomal PI(3)P homeostasis through modulation of MTM1 activity. We provide evidence that MTM1 recruitment and activity are regulated through binding to the endosomal cargo adaptor SNX17. This binding is in competition with NPxY cargo peptides, providing a molecular mechanism where cargo arrival modulates endosomal identity. Our model predicts that when cargo levels are high, SNX17 will be occupied by cargo and unable to recruit MTM1, leading to high PI(3)P levels and the establishment of the retrieval subdomain. As cargo levels decrease, SNX17 will be free to bind to MTM1, leading to PI(3)P depletion. As SNX17 is itself regulated by binding to PI(3)P, this introduces a negative feedback loop that regulates the levels of PI(3)P on endosomes in response to local cargo concentrations. Importantly, this provides a molecular mechanism to couple the membrane identity of an endosome to its functional state.

## Results

### MTM1 controls endosomal PI(3)P homeostasis

To understand the regulation of endosomal PI(3)P homeostasis, we first set out to characterise its spatial distribution in relation to retrieval subdomains that concentrate cargo for recycling back to the plasma membrane. To visualise PI(3)P, we used our recently developed recombinant biosensors [34]. This approach enables visualisation of phosphoinositides that are not effector bound after cell fixation and does not interfere with lipid signalling. Images were acquired using three colour super resolution microscopy after staining against the core Retromer component VPS35, as marker of the retrieval subdomain, together with phalloidin to detect local F-actin accumulation: another feature of the retrieval subdomain. This revealed that free PI(3)P is abundant on the bulk spherical endosomal membrane while the free level of this phosphoinositide is limited within the tubular retrieval subdomain. At the budding site, where retrieval subdomains emerge from the endosomal membrane, both PI(3)P, F-actin and VPS35 were enriched **(Fig.1a)**. To further investigate this endosomal architecture, we visualised the distribution of various PI(3)P effector proteins and the lipid kinase VPS34 together with VPS35 **(Fig. 1b)**. Both EEA1 and SNX17 followed the same distribution as free PI(3)P, as has previously been described [35]. Conversely an additional PI(3)P binding protein SNX3, that functions as a cargo-dependent membrane adaptor for the Retromer complex [36], was only present in the retrieval subdomain and not on the bulk endosomal membrane where high amounts of free PI(3)P are present. To visualise the lipid kinase VPS34, we used endogenous Halo tagged knock-in cells [37]. After inhibition of kinase activity using the small molecule VPS34-IN-1, we robustly detected VPS34 on the bulk endosomal membrane and not in the retrieval subdomain **(Fig. 1b, bottom)**. Taken together, our data reveal a domain architecture of high levels of free, not effector bound PI(3)P, generated by VPS34 on the membrane of the bulk endosomal vacuole, and a relatively low level of free, highly effector bound PI(3)P on the retrieval subdomain.

**Figure 1:**
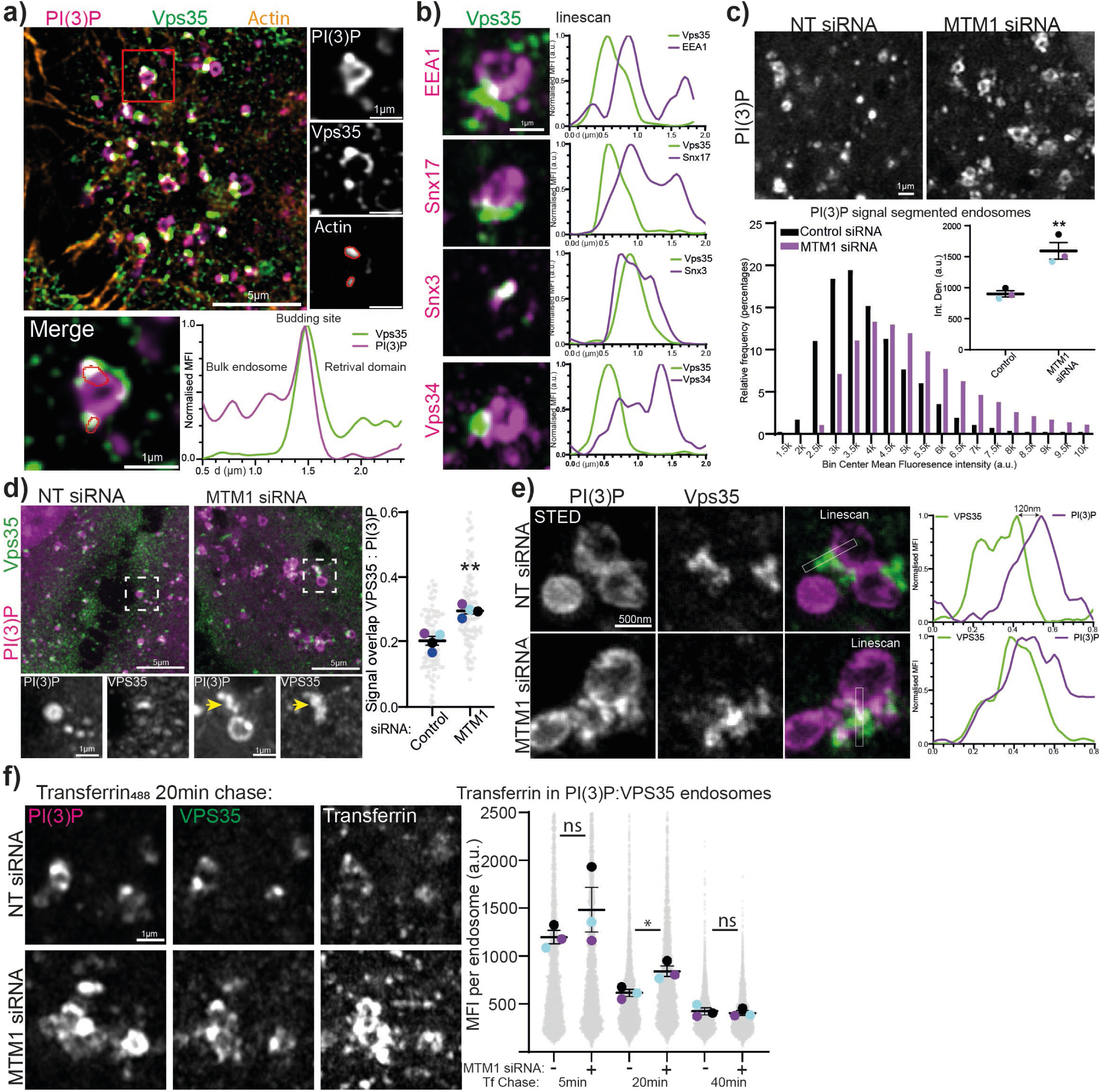
MTM1 controls endosomal PI(3)P homeostasis and segregation **a)** HeLa cells were stained against VPS35, actin and PI(3)P and imaged using structured illumination microscopy. Line scan was drawn along the retrieval subdomain and the bulk endosome membrane. **b)** HeLa Cells were stained against PI(3)P together with antibodies against SNX17, EEA1 and SNX3. HEK293T cells with endogenous Halo tagged VPS34 were treated with 1 μM VPS34-IN-1 and visualised using HaloTag Oregon green ligand. Line scans were drawn across the retrieval subdomains and the bulk endosome. **c)** HeLa cells were treated with MTM1 or non-targeting siRNA and stained against PI(3)P. Endosomal PI(3)P levels were quantified through thresholding in ImageJ with a total of 8,835 or 8,429 endosomes from three independent repeats. **d)** HeLa cells were stained against PI(3)P and VPS35 and signal overlap was analysed using the ImageJ plugin SQUASSH from 92 or 104 cells across 4 separate repeats. **e)** HeLa cells were stained against VPS35 using Alexa594 conjugated secondary antibody and PI(3)P using JF650 conjugated recombinant biosensors and image using STED microscopy. **f)** HeLa cells were treated MTM1 or non-targeting siRNA. Cells were incubated with Alexa555 labelled transferrin on ice and warmed for 5, 20 or 40 minutes to 37oC before fixation and staining against PI(3)P and VPS35. Fluorescent transferrin signal was analysed in segmented endosomes based on PI(3)P signal. Data from 18,357 segmented endosomes across different treatments and timepoints from three separate repeats.

The endosomal exit of cargo has been shown to depend on depletion of PI(3)P by MTM1[12]. To understand how MTM1 effects endosomal subdomain architecture, we depleted HeLa cells of MTM1 using siRNA and stained against PI(3)P. Consistent with previously published results [12, 13, 16, 38], we observed an elevation in endosomal PI(3)P, together with alterations in endosome morphology **(Fig. 1c).** To test the effect of MTM1 depletion on endosomal subdomain architecture we stained for VPS35 and could detect an increased amount of excess, not effector bound, free PI(3)P in the retrieval subdomain **(Fig. 1d)**. Due to the nanoscale size of endosomal subdomains, we verified these changes in PI(3)P distribution using two-colour STED microscopy **(Fig. 1e)**. To test the functional consequences of these PI(3)P changes for cargo sorting, we performed a pulse chase assay with fluorescently labelled transferrin in MTM1 depleted cells. Consistent with previous data, we detected a reduced rate of cargo recycling from endosomes, as evidenced by higher amounts of fluorescent transferrin being retained in PI(3)P and VPS35 decorated endosomes 20 minutes after ligand uptake **(Fig. 1f)**. Collectively, these data establish that MTM1 activity controls endosomal PI(3)P levels and the relative distribution of this phosphoinositide, between the bulk endosomal vacuole and the functional retrieval subdomain. A distribution control that is required for efficient endosomal cargo recycling.

Membrane recruitment regulates MTM1 activity

To gain mechanistic understanding of the regulation of MTM1 activity, we established an in vitro reconstitution system using supported lipid bilayers. While water-soluble phosphoinositide headgroups have been valuable to address substrate specificity and disease mutations, they are unable to provide detailed insights into the regulation of enzyme kinetics on membranes [39–41]. We formed supported lipid bilayers on silica beads containing 5% PI(3)P with 0.1% Atto647 labelled DOPE and added 25 nM Alexa 488-labelled recombinant probe against PI(3)P (SNAP488-2xFYVE). Confocal microscopy allowed us to directly detect PI(3)P depletion by measuring the loss of the PI(3)P probe after addition of recombinant MTM1. To quantify MTM1 activity we then used the signal of the Atto647 labelled lipids as a mask to segment individual beads to reveal the real time kinetics of MTM1 activity on membranes **(Fig. 2a)**.

**Figure 2:**
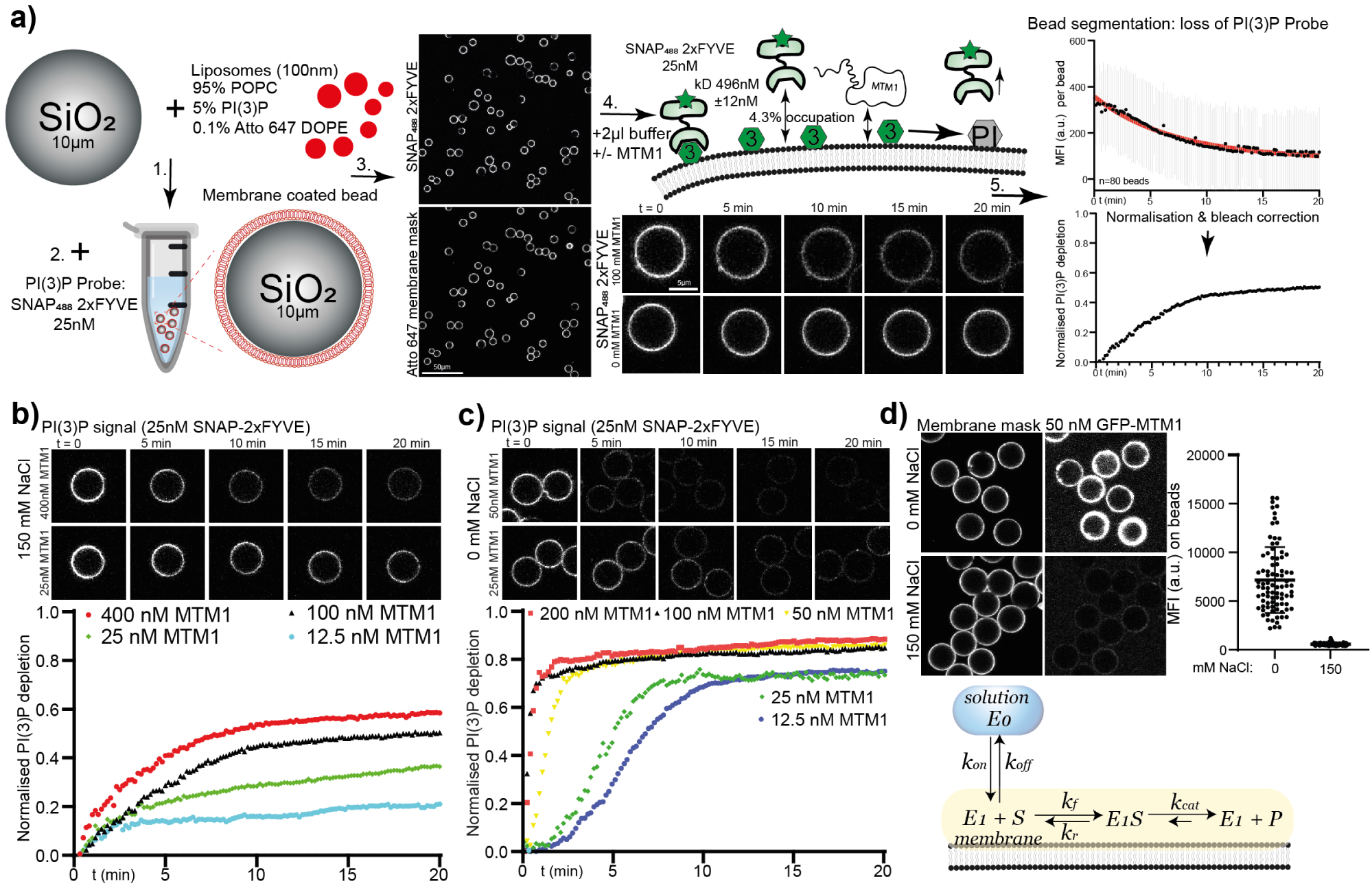
Membrane recruitment regulates MTM1 activity **a)** Overview of real time phosphoinositide conversion assay. Kinetic trace are averages from 80 individually segmented membrane coated beads using the membrane mask for thresholding. **b and c)** Kinetic traces of MTM1 activity on supported lipid bilayers at 0 or 150 mM NaCl concentrations. Each kinetic traces are averages from 59-83 individual beads per condition. **d)** GFP tagged recombinant MTM1 was added to membrane coated beads at 0 and 50 mM NaCl and membrane recruitment was analysed in ImageJ from data is Mean ±SD from 103 and 87 individually segmented beads.

Under these conditions and using physiological reaction buffers, we first noticed the relatively low activity of MTM1. Even at concentrations of 400 nM, recombinant MTM1 only led to a partial depletion of PI(3)P after 20 minutes **(Fig. 2b)**. Past studies on recombinant MTM1 have used low ionic strength buffers of <50 mM sodium chloride to assess enzyme activity. Indeed, removing sodium chloride from the reaction led to a vast increase in enzyme kinetics, with complete depletion of PI(3)P occurring within the first 2 minutes of adding 50 nM MTM1 **(Fig. 2c)**. The presence of ions typically shields electrostatic interactions, and high salt concentrations are often used to strip peripheral membrane proteins. To test if the increased enzymatic activity was due to differences in MTM1 membrane recruitment, we purified GFP-tagged MTM1 and measured its membrane binding at different sodium chloride concentrations. While at physiological salt concentrations (150 mM) almost no GFP-MTM1 signal was detectable on membranes, there was a strong enzyme recruitment in the absence of salt **(Fig. 2d)**.

The activity of membrane modifying enzymes is a combination of their affinity for the membrane and their subsequent enzymatic activity **(Fig. 2d bottom)**. The increased activity of MTM1 at low salt is therefore likely due to an increased membrane recruitment, and the low activity at physiological salt concentrations can be explained by a weak membrane binding affinity. This leads to the natural question of what mediates membrane recruitment to endosomes in the cellular environment, under physiological salt conditions?

SNX17 functions as a membrane adaptor for MTM1

To identify adaptor proteins that might mediated MTM1 recruitment to endosomal membranes we checked the BioGRID database of protein interactors [42]. Intriguingly, the endosomal cargo adaptor SNX17 has been identified as an interaction partner in five separate mass spectrometry studies [43–46], as well as one study identifying SNX27 [47]. SNX17 and SNX27 are endosomal adaptor proteins that bind to PI(3)P via a conserved PX-domain and respectively, recognise NPxY and PDZ sorting motifs in the tail of endocytic cargo proteins to promote their endosomal recycling through the retrieval subdomain. Strikingly, both sorting motifs are also present in the C-terminal tail of MTM1 and the other two class I Myotubularins, MTMR1 and MTMR2 **(Fig. 3a)**.

**Figure 3:**
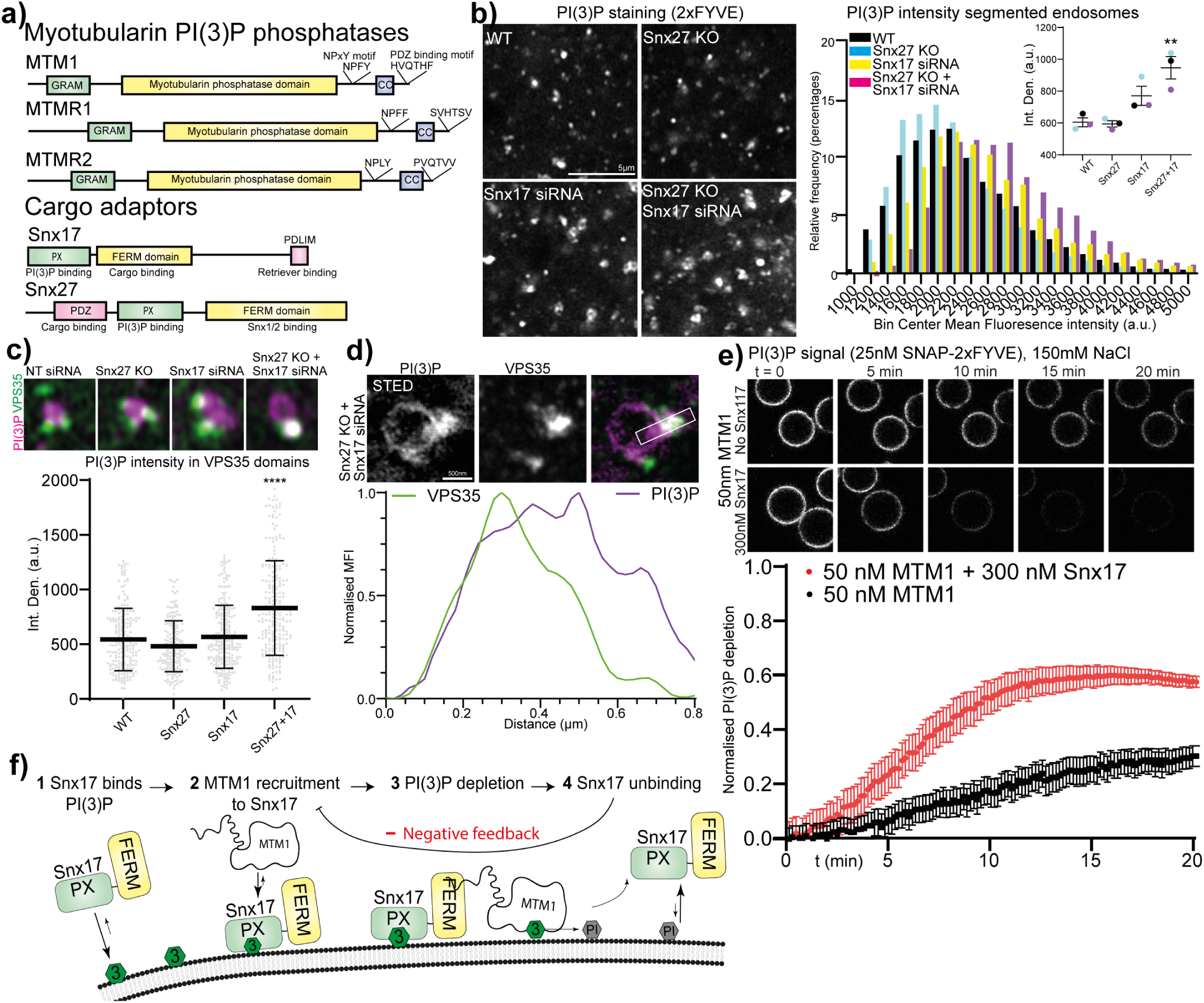
SNX17 controls PI(3)P depletion through MTM1 interaction **a)** Domain overview of class I myotubularins and the endosomal cargo adaptors SNX17 and SNX27. **b)** H4 neuroglioma cells and SNX27 knock out cells were depleted of SNX17 and stained against PI(3)P. Endosomal PI(3)P levels were quantified through thresholding in ImageJ with a total of 3,141 to 4,700 endosomes from three independent repeats. **c)** H4 neuroglioma cells and SNX27 knock out cells were depleted of SNX17 and stained against PI(3)P and VPS35. PI(3)P signal was used as mask to analyse the intensity of VPS35 signal overlapping with PI(3)P from 179 to 205 pooled from three separate repeats. **d)** H4 neuroglioma cells and SNX27 knock out cells were depleted of SNX17 and stained against VPS35 using Alexa594 conjugated secondary antibody and PI(3)P using JF650 conjugated recombinant biosensors and image using STED microscopy. **e)** Phosphoinositide conversion assay was carried out in the presence or absence of 300 nM recombinant SNX17 to measure activity of 50 nM MTM1. Kinetic traces are averages from 59 or 83 individual beads ±SD per condition. **f)** Schematic of the proposed negative feedback controlling PI(3)P depletion by MTM1 through SNX17 interaction.

If these cargo adaptors interact with MTM1 to regulate PI(3)P depletion, we would expect an increase in endosomal PI(3)P levels in their absence. To test this, we used genomic SNX27 knock-out H4 neuroglioma cells [48], together with transient depletion of SNX17 via siRNA and stained for PI(3)P. No changes in endosomal PI(3)P levels were detected in SNX27 knock-out cells. However, on its own, depletion of SNX17 led to an increased amount of endosomal PI(3)P that was further exacerbated by its depletion in the SNX27 knock-out background **(Fig. 3b)**. This suggests a level of functional redundancy between SNX17 and SNX27 in the regulation of endosomal PI(3)P homeostasis, with SNX17 being the primary player. To examine whether this increase of endosomal PI(3)P changes endosomal lipid domain segregation, we co-stained against VPS35 to reveal an accumulation of PI(3)P in the retrieval subdomain **(Fig. 3 c and d)**. Co-depletion of both SNX17 and SNX27 therefore phenocopies the depletion of MTM1.

To test if this is a direct effect of MTM1 recruitment we included recombinant SNX17 and SNX27 into our *vitro assay* to test MTM1 activity. In agreement with our cellular data, the addition of 300 nM SNX17 to membranes containing 5% PI(3)P had a strong activating effect of MTM1, leading to more than 60% PI(3)P depletion after 15 minutes **(Fig. 3 e)**. In contrast, addition of SNX27 had only minor effects on MTM1 activity and at high concentrations blocked phosphoinositides conversion, possibly through competitive binding with PI(3)P **(Fig S. 1 a)**. Importantly, because SNX17 is recruited to membranes through binding to PI(3)P via its PX domain, its recruitment of MTM1 introduces a negative feedback loop, in which the activity of MTM1 will lead to the displacement of its own membrane adaptor and thereby negatively regulate its own activity **(Fig. 3f).**

To directly test this model, we transfected HeLa cells with GFP tagged wild type and catalytic dead (C375S) MTM1 and assessed their subcellular localisation. In line with its low membrane binding affinity, both constructs were completely cytosolic with limited detectable endosomal association **(Fig 4a left)**. However, upon co-expression of HA tagged SNX17 we observed robust endosomal localisation of MTM1-C375S but not of wild type MTM1 **(Fig 4a right)**. This provides further evidence that MTM1 is recruited to membranes by SNX17 and that its activity negatively regulates this recruitment.

**Figure 4:**
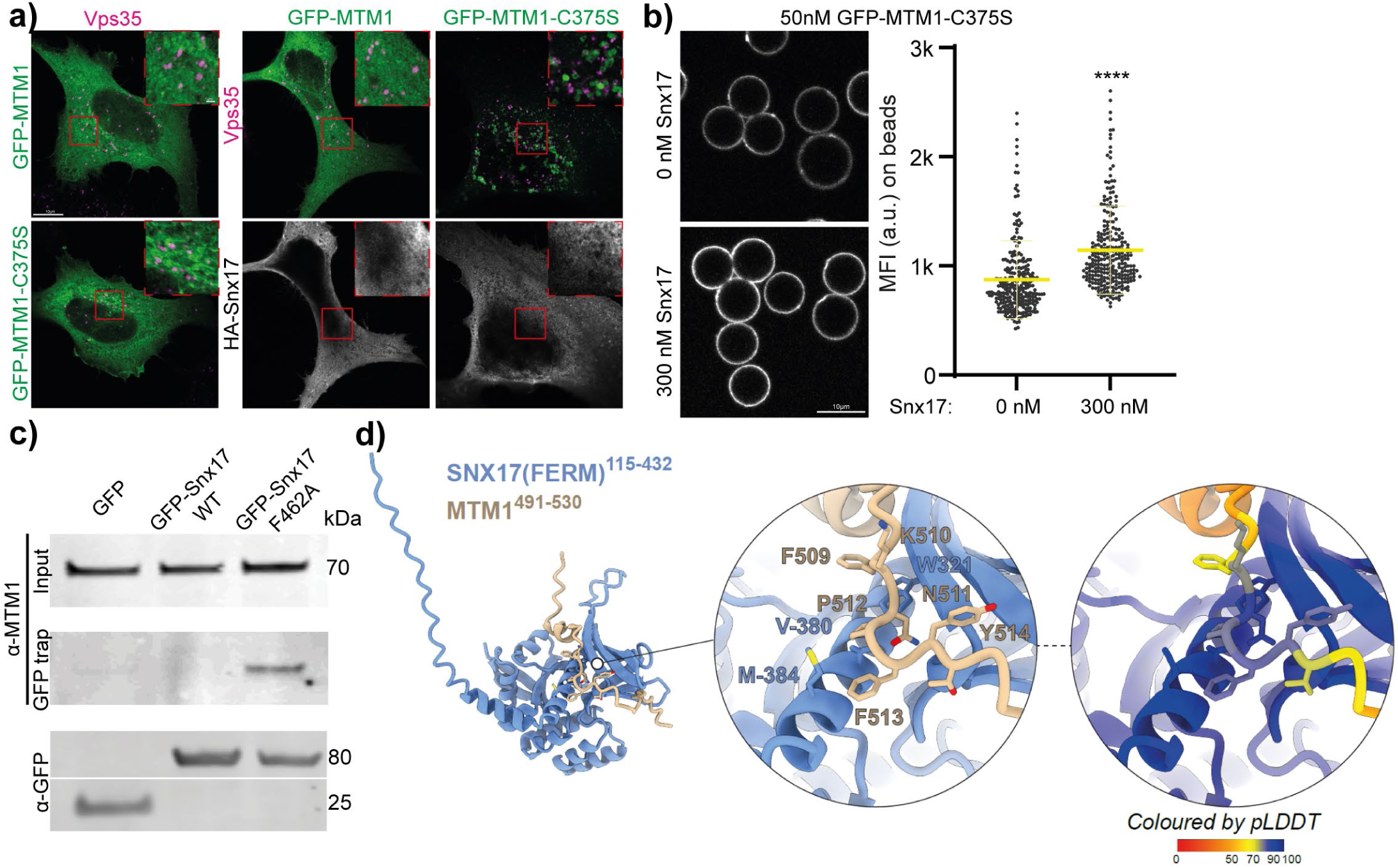
MTM1 binds to the cargo binding domain of SNX17 for membrane recruitment **a)** HeLa cells were transfected with GFP-MTM1 or catalytic dead GFP-MTM1-C375S alone or in combination with HA tagged SNX17 and stained against VPS35 and the HA tag. **b)** Membrane recruitment to membrane coated beads of recombinant GFP tagged MTM1-C375S was measured in the absence and presence of 300 nM recombinant SNX17. n=266 and 274 individually segmented beads. **c)** HeLa cells were transfected with constructs for expression of GFP, GFP-SNX17 or GFP-SNX17-F462A. Co-IP was carried out using GFP trap agarose followed by western blot with MTM1 and GFP antibodies. **e)** Alphafold model of MTM1 region 491-530 containing with the 509-FKNPFY-514 motif engaging with the canonical cargo binding groove of the SNX17 FERM domain.

To directly validate that SNX17 is a membrane adaptor for MTM1, we assessed membrane recruitment of recombinant GFP tagged MTM1-C375S to supported lipid bilayers. Addition of recombinant SNX17 led to a significant increase in the recruitment of catalytically dead MTM1 to membranes containing 5% PI(3)P **(Fig. 4b)**. To better understand this interaction, we made use of a specific point mutation in SNX17. In its basal state SNX17 is partially autoinhibited through its C-terminus folding into the cargo binding pocket in its FERM domain and this autoinhibition can be released by a single F462A mutation [35, 49]. Pulldowns of cells transfected with GFP tagged wild-type SNX17 or SNX17-F462A showed a specific interaction of endogenous MTM1 with the autoinhibited released mutant of SNX17 **(Fig. 4c)**. When we co-expressed this SNX17 mutant together with GFP tagged wild type or catalytic dead MTM1, we observed the formation of large cytosolic aggregates containing both SNX17 and MTM1-C375S **(Fig. S1d)**. To test if the NPxY motif in the C-terminal tail of MTM1 is a suitable ligand for SNX17 we used AlphaFold modelling of amino acids 491-530 in MTM1 together with the FERM domain of SNX17. This predicts that the region 509-FK<u>NPFY</u>-514, containing the NPxY motif, folds into the cargo binding domain of SNX17 with high confidence **(Fig. 4d)**. Taken together this data suggests that the NPxY motif in MTM1 functions as a ligand for Snx17 and that this interaction drives MTM1 membrane recruited and thereby activity.

### Cargo regulates PI(3)P depletion through competition with MTM1 for SNX17 binding

If MTM1 and cargo contain the same sorting motifs and share the same binding site on SNX17, then membrane localised cargo should be in competition with MTM1 for SNX17 binding. High cargo levels should outcompete MTM1 and thereby block the activating effect of SNX17 on PI(3)P depletion. To directly test this, we modified our *in vitro* phosphoinositide conversion assay to include membrane bound cargo peptides. We prepared two separate populations of membrane coated beads containing either 0.1% Atto647 or Rhodamine DOPE fluorescent lipids together with 5% PI(3)P. To the beads with Rhodamine DOPE we used click chemistry to attach the SNX17 cargo peptide of LRP1, and to the other beads a mutant peptide, in which the SNX17-binding NPxY motif was replaced with alanine (AAAA). We then mixed both populations, added 300 nM SNX17 and started the reaction by addition of 50 nM MTM1. We segmented the separate populations based on their fluorescent lipids and analysed the kinetics of MTM1 activity on each bead population within the same reaction volume **(Fig. 5a)**. The activity of MTM1 was markedly decreased (by about 50%) on membranes containing the LRP1 cargo peptide, while PI(3)P was rapidly depleted on membranes containing the AAAA mutant peptide. Crucially, these data directly link cargo concentration to membrane identity conversion through competitive binding of MTM1 and cargo to SNX17.

**Figure 5:**
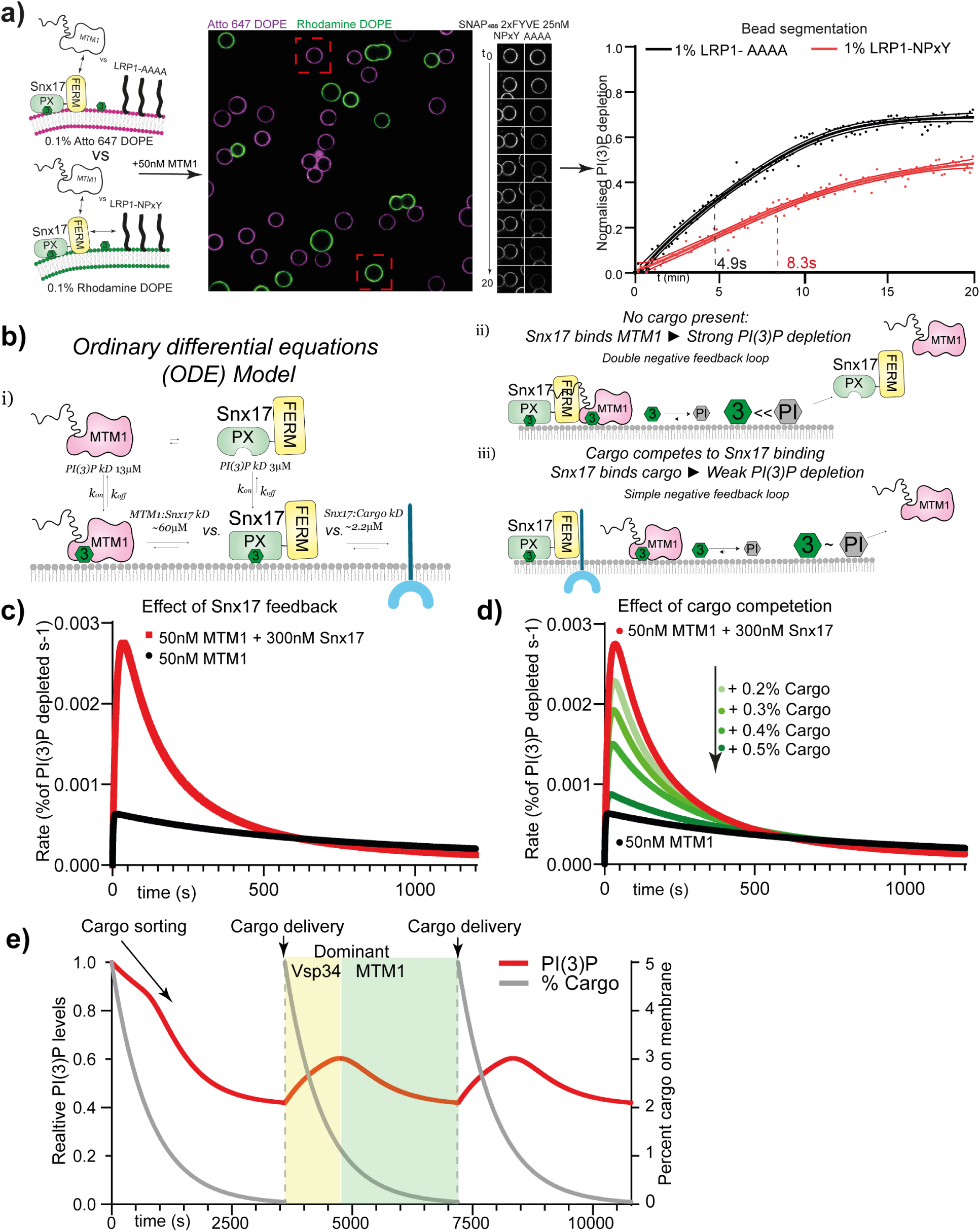
Cargo competes with MTM1 binding to SNX17 to regulate PI(3)P depletion **a)** Phosphoinositide conversion assay on membrane coated beads containing either LRP1_NPxY or LRP1_AAAA cargo peptides. Data is average from 16 or 28 individually segmented beads fitted with a five-parameter logistic equation and 99% confidence interval. **b)** Schematic of the Ordinary Differential Equations (ODE) model to determine the effect of negative feedback and cargo competition on the rate of PI(3)P depletion by MTM1. **c)** Results of the ODE model of the rate of PI(3)P depletion by MTM1 in the presence or absence of 300 nM SNX17. **d)** Results of the ODE model in the presence of increasing amount of membrane localised cargo on the rate of PI(3)P depletion by MTM1. **e)** Results of the ODE model on the relative PI(3)P levels including VPS34 mediated PI(3)P generation and changing cargo concentrations

Our data so far indicates that SNX17 is a membrane adaptor for MTM1 and that this interaction is in competition with cargo binding. SNX17 is itself recruited to membranes by binding to PI(3)P, introducing negative feedback into the reaction. To better understand how this negative feedback and cargo occupancy regulate endosomal PI(3)P homeostasis we established an ordinary differential equations (ODE) model (see Materials and Methods). This model allows us to calculate the effect of SNX17 binding, negative feedback and cargo competition on the rate of PI(3)P depletion by MTM1 **(Fig. 5b)**.

In the absence of cargo, the addition of SNX17 into the ODE model leads to a rapid, but transient, increase in the rate of PI(3)P depletion through increasing the dwell time of MTM1 on the membrane. However, as MTM1 depletes PI(3)P, SNX17 is lost with a concomitant decrease in the recruitment of MTM1 and therefore the rate of phosphoinositide conversion **(Fig. 5c)**. The spike followed by a steep decline in the rate of PI(3)P depletion is the consequence of the negative feedback. When we include increasing amounts of membrane localised cargo to the model, we see a corresponding dampening of this activating effect of SNX17 **(Fig. 5d)**. We then expanded our model to include generation of PI(3)P to represent VPS34 activity, including negative feedback by high PI(3)P levels as shown by [50, 51] and outlined in the Discussion. We then added repeating spikes of cargo to the model, followed by its exponential decay, to simulate cargo delivery and sorting **(Fig. 5e)**. The initial state of high PI(3)P is followed by its decay due to the negative feedback in VPS34 activity. As cargo levels drop, SNX17 becomes available for MTM1 binding, leading to an increased PI(3)P depletion until the negative feedback balances PI(3)P depletion with VPS34 mediated PI(3)P generation. Adding cargo back into the system then leads to an increase in PI(3)P due to SNX17 being cargo bound and decreasing MTM1 activity with VPS34 being dominant. Then, as PI(3)P increases, more SNX17 is recruited to the membrane that is initially occupied by cargo until the cargo levels drop again for MTM1 to engage SNX17 and to become dominant over Vps34 activity **(Fig. 5e)**. This inverse regulation of MTM1 and VPS34, through a cargo sensitive feedback loop, can therefore adapt endosomal PI(3)P levels in response to cargo sorting and delivery.

Under homeostatic conditions the amount of cargo entering sorting endosomes via fusion of early endosomes will be in constant flux with cargo leaving the system through budding from retrieval subdomains and intra-luminal vesicles from the degradative subdomain. To disrupt this equilibrium, we treated cells with hypertonic media to completely block endocytic cargo uptake from the plasma membrane [52]. In line with our model, this acute block led to a rapid drop in endosomal PI(3)P levels, as seen both in fixed cells after 30 minutes and in live cell microscopy of cells expressing the PI(3)P biosensor **(Fig. 6a and S2 a)**. Inversely, to block cargo exit, we blocked branched actin formation with the ARP2/3 complex inhibitor, CK666. This blocks the formation of retrieval subdomains, leading to enlarged swollen endosomes. These enlarged endosomes also contained an increased total amount of PI(3)P **(Fig. 6b)**, strengthening our mechanistic link between endosomal cargo concentrations and PI(3)P levels.

**Figure 6:**
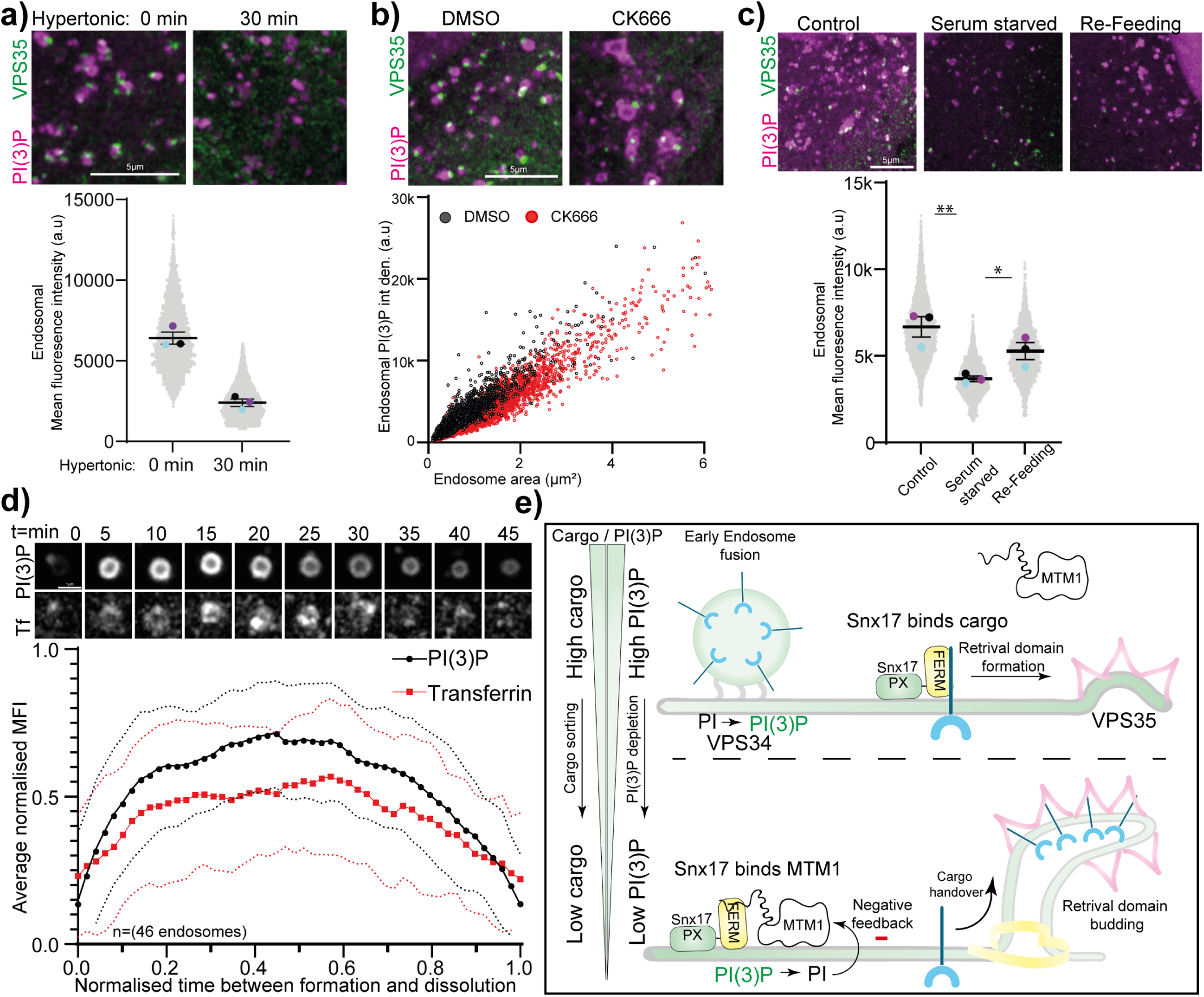
PI(3)P levels are linked to the cargo levels of endosomes. **a)** HeLa cells were treated with hypertonic media for 30 minutes and stained against PI(3)P and VPS35. Endosomal PI(3)P levels were quantified through thresholding in ImageJ with a total of 3,475 or 8,461 endosomes from three independent repeats. **b)** HeLa cells were treated with 10 μM CK666 for 1 hour and stained for PI(3)P and VPS35. Endosomal PI(3)P levels and size were quantified through thresholding in ImageJ with a total of 19,006 or 26,310 endosomes from three independent repeats. **c)** HeLa cells were serum starved for 2.5 hours and fixed directly or refed with complete media for 2.5 hours and then fixed and stained against PI(3)P and VPS35. Endosomal PI(3)P levels were quantified through thresholding in ImageJ with a total of 491 to 1,529 endosomes from three independent repeats. **d)** HeLa cells expressing 2xFYVE-eGFP were serum starved for 3 hours before being imaged live at 37°C in CO2 full media containing together with 5 μg/ml Alexa647 labelled transferrin for 2 hours with one complete Z-Volume captured every minute. 46 individual endosomes were manually tracked, and fluorescents intensities of PI(3)P and transferrin were normalised, averaged ±SD and linearly resampled to normalise the time between formation and dissolution of each endosome. **e)** Model of cargo dependent PI(3)P regulation through direct feedback between SNX17 and MTM1. At high cargo levels SNX17 is occupied by cargo, and retrieval subdomains are formed, followed by cargo sorting. At low cargo levels SNX17 is free to bind to MTM1, leading to PI(3)P depletion and endosome maturation.

Both of these treatments are strong perturbations that heavily alter signalling cascades and cellular functions. To test more physiological conditions, we used serum starvation, reasoning that this should lead to reduced receptor engagement and recycling. Indeed, starvation for 2 hours led to a drastic decrease in endosomal PI(3)P levels, and the amount of PI(3)P positive endosomes per cell **(Fig S2 b)**; this recovers nearly back to baseline after refeeding with serum for 2 hours **(Fig 6c)**. The recovery of endosomal PI(3)P upon refeeding indicates the formation of new endosomes as cargo uptake is restored. To directly visualise this *de novo* formation, we starved cells expressing the PI(3)P biosensor for 3 hours and then started live cell imaging of whole Z-volumes after addition of serum together with fluorescent transferrin as a tracer of endocytic uptake and recycling. Manual tracking enabled us to detect multiple examples of the *de novo* formation of long-lived sorting endosomes, with a lifetime of 10 to 50 minutes, that increased in PI(3)P intensity together with transferrin uptake **(Fig 6d)**. Importantly, sorting and decrease in cargo was associated with a concomitant drop in PI(3)P intensity. These data directly support our link between endosomal cargo levels and local PI(3)P regulation.

Taken together our data supports a model that tightly links endosomal cargo levels to PI(3)P homeostasis: when cargo levels are high, SNX17 is bound to cargo, PI(3)P depletion is blocked, and PI(3)P is available for formation of the SNX3-Retromer-containing retrieval subdomain. As cargo is sorted away from the bulk endosomal membrane, MTM1 will be able to bind to SNX17 and depletes PI(3)P. This depletion starts a negative feedback loop through the concomitant loss of SNX17 and thereby links PI(3)P depletion to endosomal cargo levels. **(Fig. 6e)**. Importantly, this mechanism links endosomal membrane identity with its functional state in sequence-dependent cargo recycling through the retrieval subdomain.

## Discussion

Cargo delivery and exit from endosomes are dependent on effector proteins that bind to PI(3)P. When PI(3)P levels are either too high or too low cargo fails to be sorted correctly with catastrophic consequences to cellular function. As is the case in X-linked centronuclear myopathy (XLCNM). Here we have sought to understand how PI(3)P homeostasis is regulated and how this is coupled to the function of endosomes in cargo sorting.

Depletion of MTM1 leads to the well described increase in endosomal PI(3)P levels and impairment of cargo recycling [12, 13, 16, 38]. Additionally, we could observe a breakdown of lipid domain segregation. Understanding endosomal PI(3)P homeostasis therefore requires understanding of MTM1 regulation. Through a combination of *in vitro* reconstitution, cell biology and computational modelling we have determined that PI(3)P depletion by MTM1 is regulated through membrane recruitment by binding to the endosomal cargo adaptor SNX17. This interaction is mediated by sorting motifs in MTM1 and in competition with the same sorting motifs in cargo that are recognised by SNX17. It has been well established that these motifs in cargo direct their endosomal sorting through engaging with adaptor proteins [29, 35]. Intriguingly, the presence of the same motifs in the C-Terminal tail of cytosolic MTM1 allows the underlying phosphoinositide identity of the membrane to be linked to the endosomal sorting of cargo. Importantly, this mechanism couples endosomal cargo levels to PI(3)P depletion. Meaning that when cargo is high, PI(3)P is high and *vice versa*. A connection that we directly observed on newly formed endosomes.

Endosomes are highly dynamic organelles that change phosphoinositide identity through MTM1 recruitment depending on the metabolic state of the cell, such as starvation. During which endocytic receptor recycling will be reduced, leading to a low endosomal cargo density, which would explain the increased MTM1 activity seen under these conditions [17, 18, 53]. Our proposed model is therefore able to provide mechanistic detail of how endosomes change membrane identity depending on their functional state - SNX17 may function not only as a cargo adaptor but as a molecular sensor that couples cargo occupancy to endosomal membrane identity. Thus, as cargo is shuttled into retrieval domains [31, 35, 49] and its concentration within the bulk endosomal membrane decreases, SNX17 becomes free to engage MTM1 to reset the PI(3)P levels. Then as new cargo is being delivered, SNX17 will be bound to cargo, PI(3)P levels will increase, retrieval subdomain will form and the cycle continues.

The regulation of lipid kinases and phosphatases through binding to membrane adaptors is a common theme and allows for the establishment of feedback loops that couple enzyme activity to the levels of its substrate and/or product [19, 20, 22, 23, 54, 55]. As SNX17 is recruited to membranes in part by PI(3)P binding [8], this introduces negative feedback into the regulation of MTM1 and couples its activity to the availability of its substrate. Importantly, this depletion of PI(3)P is balanced with its generation by the lipid kinase VPS34. The observation that endogenously tagged VPS34 shows increased membrane localisation after inhibition of its catalytic activity [37], suggests that its activity also negatively regulates its membrane recruitment. A potential mechanism being that VPS34 is recruited to membranes through binding to active Rab5 [19, 20] and that TBC-2 (a Rab5-GAP) is recruited to endosomes through PI(3)P to inactivate Rab5 in C-elegans [50]. This model would imply that high PI(3)P levels both negatively regulate VPS34 through Rab5 inactivation, as well as activating MTM1 through SNX17 recruitment. This inverse regulation of MTM1 and VPS34 activity through PI(3)P sensitive feedback loops has the potential to balance endosomal PI(3)P levels.

Finally, it is likely that there are multiple additional layers for regulation of endosomal PI(3)P homeostasis. MTM1 is known to form both homodimers as well as heterodimers with catalytical dead MTMR12 which is supposed to have a regulatory effect on its activity [56, 57]. It is also possible that MTM1 undergoes membrane mediated dimerisation after having been enriched at the membrane, as is the case for PIP5K [58]. Additionally, post translational modifications have been shown to regulate membrane recruitment of the related MTMR2 protein to the Commander complex on both endosomes and phagosomes [59–61]. Future work will have to address how dimerisation and/or post-translational modifications might regulate MTM1 activity and if there are similar regulatory feedback loops in the control of the wider MTM family.

## Acknowledgements

1. H. Maib is supported by a Wellcome Trust early Career award 225528/Z/22/Z. Anwar Sadat is supported by a Wellcome Trust Discovery Award 311591/Z/24/Z. We acknowledge the Wolfson Light Microscopy Facility for the use of the LSM 980 Airyscan 2 supported by the Medical Research Council grant MR/X012077/1. We further thank Prof. Jason King and Prof Elizabeth Smythe for their helpful comments on the manuscript.

Research in the Cullen lab is supported by the Wellcome Trust (220260/Z/20/Z and 319367/Z/24/Z), Medical Research Council (MR/P018807/1), Lister Institute of Preventive Medicine, and through award of a Royal Society Research Professorship (RSRP/R1/211004 and RSRPR26/1003).

## Materials and Methods

### Cell culture and transfection

All cell lines were cultured in DMEM supplemented with 10% FCS and penicillin/streptomycin and passaged every 2–3 days. The HeLa cell line was a kind gift from Prof. Elizabeth Smythe (University of Sheffield, Sheffield, UK). The SNX27 knock out H4 neuroglioma cells were generated previously [48]. The endogenous Halo-VPS34 HEK293T [37] cells were a kind gift from Prof. Dario Alessi (University of Dundee, UK). For siRNA mediated knockdown of MTM1 and SNX17, cells were grown on 24 mm 1.5 borosilicate coverslips to a confluency of ∼30-50% and transfected with a SMARTpool of 5 siRNAs against MTM or SNX17 (Horizon Discovery/Dharmacon) DharmaFECT 1 Transfection reagent. To ensure optimal knock down, cells were transfected on two consecutive days. For transient overexpression of GFP tagged MTM1, MTM1-C375S, 2xFYVE, HA-SNX17 and HA-SNX17-F462A, cells were transfected using PolyFect Transfection Reagent (Qiagen) and imaged the next day.

### Cell staining

Cells were stained against PI(3)P using recombinant biosensors as described in [34]. Briefly, the biosensor against PI(3)P was expressed in BL21, purified via the N-Terminal 6xHis tag by three step purification of Ni^2+^ affinity, ion exchange chromatography followed by gel filtration. The recombinant biosensor was labelled in vitro with SNAP-Surface Alexa Fluor 647 (NEB) and stored in 5 µM aliquots at-70°C. For staining of cells using the recombinant biosensor, cells were fixed with 4% PFA with 0.2% glutaraldehyde (EM grade) warmed up to 37°C for 20 minutes followed by two quick washes and incubation with 50 mM NH4Cl in PBS for 20 minutes. Coverslips were transferred onto ice, and the cells were washed with ice-cold PIPES buffer. Cells were stained, blocked, and permeabilized at the same time in 5% BSA + 0.5% (wt/vol) Saponin (S7900 from quillaja bark; Sigma-Aldrich) in PIPES with the addition of labelled biosensors to a final concentration of 500 nM and together with Goat polyclonal antibody to VPS35 (Antibobies.com A83699) for 60 minutes on ice. Cells were washed three times in ice cold PIPES, incubated with fluorescently labelled secondary anti goat antibodies for 45 minutes and washed three times with ice-cold PIPES. Cells were post-fixed with 2% PFA in PBS on ice for 10 minutes before being returned to room temperature for another 10 minutes. Cells were washed with 50 mM NH4Cl in PBS at room temperature, rinsed with ddH2O, mounted, and sealed using Prolong Gold.

For co-staining of VPS35 together with EEA1 (mouse ab70521; Abcam), SNX17 (rabbit HPA043867, Sigma-Aldrich) and SNX3 (Proteintech mouse monoclonal 2B5B7), cells were fixed in 4% PFA for 20 minutes, followed by quenching in 50 mM NH4Cl in PBS for 20 minutes. Cells were permeabilized using 0.1% Triton X-100 and blocked with 1% BSA for 1 hour, stained with primary antibodies for 1 hour in PBS, washed with PBS, stained with fluorescently labelled secondary antibodies, washed again and sealed and mounted using prolong gold mounting media. Endogenous Halo tagged VPS34 was visualised after treatment with DMSO or VPS34-IN-1 (MedChemExpress) at 1 µM for 30 minutes using the Halo-OregonGreen (Promega) at 250 nM at 37°C while live. Cells were then fixed and stained against VPS35 as described above.

For Transferrin pulse chase assay, HeLa cells were grown on coverslips and chilled on ice for 10 minutes in ice-cold PBS Transferrin conjugated to Alexa568 (Thermo Fisher Scientific) was bound to the cell surface at 5 μg/ml for 30 minutes in PBS on ice followed by three wash steps on ice. Cells were transferred into prewarmed (37°C) full growth media for indicated timepoints before being fixed and stained against PI(3)P and VPS35 as described above.

### Microscopy

Confocal imaging was carried out in Z-Stacks of the mid-section of cells 150 nm step size using LSM980 Airyscan with a Plan Apochromat 63 × 1.4NA lens in SR mode using full 32 detector array to capture maximum spatial information and processed for 3D Airyscan processing.

STED imaging was performed on a commercial expert line Abberior STED microscope equipped with 485, 561, and 645 nm excitation lasers and an Olympus Objective UPlanSApo 100× 1.40 NA. For two-color STED of Vps35 and PI(3)P, cells were stained as described before with biosensors against PI(3)P conjugated to SNAP ligand JF650 and secondary antibodies conjugated to Atto594. The 775-nm depletion laser was used to deplete dyes in two-color STED experiments. Two-color images were recorded sequentially line by line. Detection was carried out with avalanche photodiodes with detection windows set to 571–630 nm and 650–756 nm. The acquisition was carried out with Instruments Development Team, Imspector Image Acquisition & Analysis Software v16.3.16118 (https://www.imspector.de). The pixel size was set to 20 nm, and Raw STED images were deconvolved to reduce noise using Richardson–Lucy deconvolution from the Python microscopy PYME package (https://python-microscopy.org/).

For 3D structured illumination microscopy, cells were stained with recombinant biosensors against PI(3) conjugated to Alexa647, together with Vps35 staining in Alexa488 and Phalloidin Texas Red (ThermoFisher). Cells were imaged using the ZEISS Elyra 7 microscope operating in Lattice SIM mode through Plan-Apochromat 63x/1.40 NA oil-immersion objective lens. Excitation was performed using 488, 561, and 640 nm laser lines. Fluorescence was detected using Hamamatsu ORCA-Flash4.0 cameras and a DuoLink SR QUAD filter module equipped with two filter cubes (LP 560 and BP 490–560/LP 640) and corresponding emission filters for multichannel fluorescence detection. SIM images were acquired as z-stacks comprising 27 planes with a z-step of 120 nm using a 25 ms exposure time. Image acquisition and SIM processing was controlled using ZEN software, with channel alignment applied to the multichannel images.

### Protein Purification

Recombinant proteins were expressed in BL21 Rosetta(DE3) E. coli cells. Four litres of cell culture was grown in baffled Erlenmeyer flasks using a shaking incubator (37°C; 180 RPM) until an OD of 0.6 - 0.8 was reached. Protein expression was induced with 0.5 mM isopropyl β-D-1-thiogalactopyranoside (IPTG), and the cell culture was left in a shaking incubator overnight at a lower temperature (16°C; 180 RPM). Cells were spun at 5,000 RCF for 20 minutes (4°C). The cell pellet was resuspended to a final volume of 30 mL using 4°C standard buffer (20 mM HEPES, 150 mM NaCl, 0.1% TCEP, pH 7.4). Three complete Mini EDTA-free protease inhibitor tablets (Sigma-Aldrich) were added and Resuspended pellets were lysed with sonication and spun at 55,000 RCF for 45 minutes (4°C). The lysate was pushed through a 0.45 μm Minisart filter (Sartorius) to remove cellular debris.

GFP-tagged MTM1 and MTM1-C375S were purified via an N-Terminal TwinStrep tag using a 5 mL StrepTrap XT column (Cytiva) using an ÄKTA Pure system (Cytiva). The Streptavidin column was washed with 75 mL (15 column volumes) of standard buffer. MTM1 was eluted isocratically using 15 mL standard buffer containing 50 mM of biotin. For in vitro reconstitution experiments the recombinant PI(3)P biosensor was expressed with an N-terminal a 6xHistidine tag. The lysate was run through a 5 mL HisTrap HP column (Cytiva) and washed with 100 mL (20 column volumes) of standard buffer. The probe was eluted using a gradient of standard buffer containing 250 mM of imidazole (from 0% - 100 % elution buffer).

Wild type MTM1, SNX17 and SNX27 were purified using an N-Terminal GST tag followed by an HRV 3C cleavage site. Lysate was incubated with Glutathione Sepharose beads (Protein Arc) for 2 hours on a rocker at 4°C. The beads were then washed five times with 10 mL of standard buffer before being resuspended in 5 mL standard buffer. The beads were then incubated overnight on a rocker at 4°C with PreScission Protease. Beads were spun at 1000 x RCF for 2 minutes and the supernatant containing cleaved MTM1 was collected. All proteins were purified using a finale gel filtration step on either an HiLoad 16/60 Superdex 200 (Cytiva) or Superdex 200 10/300 GL column (Cytiva). Fractions containing purified protein were pooled and concentrated using a Vivaspin 20 concentrating (Sartorius) and aliquots and stored at-80oC following snap-freezing.

### SNAP-labelling

The recombinant PI(3)P biosensor contains an N-Terminal SNAP tag and labelling was performed mixing 5 μM of protein with 10 μM SNAP-Surface Alexa Fluor 488 dye (New England BioLabs) and incubated for 120 minutes in the dark at 4°C. Free dye was removed using a Vivaspin 20 concentrating column (Sartorius). 20 mL of standard buffer (20 mM HEPES, 150 mM NaCl, 0.1% TCEP, pH 7.4) was passed through the concentrator at 4°C until a final volume of 100-200 µL was achieved. The labelling efficiency was determined by measuring absorbance at A280 absorption and A488, snap-frozen at-80oC for storage.

### Membrane coated bead preparation

Membrane-coated beads were prepared using the method as described in (Maib and Murray, 2022). Briefly, liposomes were produced by mixing 5 mol % phosphatidylinositol 3-phosphate diC16 (Echelon) with 95 mol % 1-palmitoyl-2-oleoyl-sn-glycero-3-phosphocholine (Avanti) and 0.1% Atto647N-DOPE (Avanti).Mixtures were evaporated using Nitrogen gas to remove chloroform and resuspended in 1 mL standard buffer (20 mM HEPES, 150 mM NaCl, 0.1% TCEP, pH 7.4) and subjected to six freeze-thaw cycles using liquid nitrogen and a water bath heated at 37°C. Lipid suspensions were then passed through a 100 nm filter (Whatman Nuclepore) 11 times to produce liposomes of approximately 100 nm in diameter. Liposomes were stored at 4°C and used within 24 hours. Membrane-coated beads were generated by mixing 10 µg of liposomes with ∼0.5 × 10^6^ 10 µm silica beads (Whitehouse Scientific) in a final volume of 100 μL and NaCl concentration of 200 mM. The mixture was left on a rotor for 30 minutes at room temperature. Beads were then washed twice in 20 mM HEPES, once in buffer containing 20 mM HEPES, 150 mM NaCl, and 0.1% TECEP and one final time in buffer containing 20 mM HEPES, 150 mM NaCl, 0.1% TECEP, and 2 μg/mL Casein for blocking of unspecific binding.

### Phosphoinositide conversion assay

Kinetic experiments were prepared by mixing 10 μL of freshly prepared Membrane-coated beads up to 18 μL with buffer/protein (e.g. SNX17), 488-labelled PI(3)P probe (final concentration of 25 nM) and buffer (20 mM HEPES pH 7.0, 150 mM NaCl, 0.1% TECEP, and 2 μg/mL Casein). The 18 μL bead solution was added to the corner of an uncoated µ-Slide 8-well chamber (Ibidi). Components were allowed to equilibrate for 5-10 minutes before the slide was mounted on an LSM980 Airyscan 2 microscope (Zeiss) using a 64x oil objective. 2 μL of reaction buffer containing recombinant MTM1 was added to 18 μL of beads directly proceeding imaging. Beads were imaged with single confocal slices at the equatorial plane every 10-20 seconds for of 10-30 minutes.

Loss of recombinant PI(3)P biosensor was analysed using a custom ImageJ script that segments the fluorescence channels based on the Atto647N-DOPE signal as membrane mask. The mean fluorescence intensity (MFI) of each imaging frame was normalised against the starting intensity at t=0. This resulted in a normalised plot of % PI3P depletion over time. Note that only ∼4% of PI(3)P molecules were masked by PI(3)P probe binding during initial assay conditions meaning that competition with MTM1 in binding PI(3)P was not major factor.

For cargo competition assays, two separate membrane coated bead populations were prepared from liposomes containing 5% PI(3)P and 1% DBCO-PE, and 0.1% Atto647N-DOPE or 5% PI(3)P and 1% DBCO-PE, and 0.1% Rhodamine DOPE. Peptides were attached to the membranes using copper free click chemistry via the DBCO-PE and N-Terminal Azide residues on the peptides: (Lys(N3)GCGSGSGSRMTNGAMNVEIGNPTYKMYEGGEPDDG(Coumarin}) LRP1:AAAA (Lys(N3)GCGSGSGSRMTNGAMNVEIGAAAAKMYEGGEPDDG(Coumarin)) at a ratio of DBCO:peptide 1:3 at 4°C in standard buffer overnight.

### Determination of binding affinity of recombinant PI(3)P biosensor

488-labbled PIi(3)P probe was titrated in a dilution series from 1500 nM to 23 nM with a final buffer composition of 20 mM HEPES pH 7.0, 150 mM NaCl, 0.1% TECEP, and 0.2 mg/mL Casein. 10 μL of each Pi(3)P titration was mixed with 10 μL of freshly prepared Membrane-coated beads. Beads were allowed to equilibrate at room temperature for 20 minutes before imaging on LSM980 Airyscan 2 microscope.

The mean 488 fluoresce on membrane coated beads at each concentration was plotted to yield a binding curve against the constant concentration of Pi(3)P as calculated at 110 nM (see below) (Fig. S2). Non-linear regression using a mass-action equation shown below produced a binding affinity of 457 nM, translating to 4.3% of PI(3)P molecules are bound/masked by the probe during initial assay conditions.

The concentration of PI(3)P molecules was calculated the using the method described in (Hansen, et al. 2019). Briefly, the number of beads in each reaction volume was counted using an Olympus CK40-SLP microscope (Olympus) and counting chamber (Hirschmann). Assuming the mean radii of beads was 5 µM, this gives a mean surface area of 314 µm^2^ per bead. The surface area of a single DOPC headgroup was 7.2 x 10^-7^ µm^2^. Given this, the number of beads in an assay, and the fact that PI(3)P made up 5% the total mol% of lipids, the molar concentration of PI(3)P in each assay could be determined by counting the beads beforehand.

### SNX17-bead affinity measurement

The affinity of recombionant SNX17 to PI(3)P was determined by a competetion assay against recombinant PI(3)P biosensor. Fresh membrane-coated beads were prepared and incubated for 20 minutes at room temperature with 100 nM 488-labbeled PI(3)P probe together with either

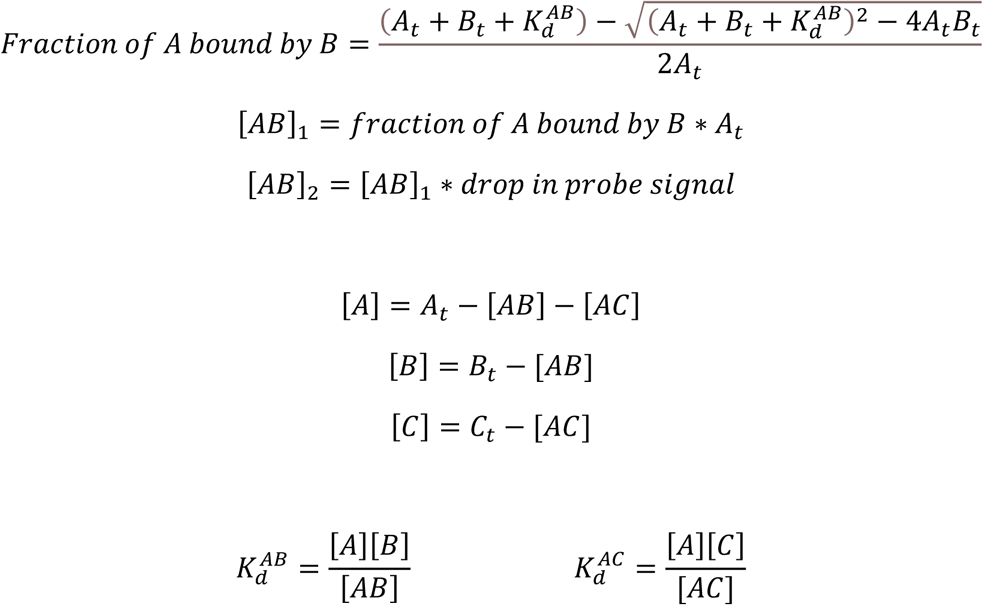

1500 nM, 750 nM, 375 nM, or 0 nM SNX17. A linear decrease in PI(3)P probe binding was observed alongside increasing concentration of SNX17 demonstrating competition for binding PI(3)P (Fig. S1b). Given the following mass action equations, the SNX17:PI(3)P affinity was 3.16 µM.

### Ordinary differential equation (ODE) modelling

The ODE models (MTM1_ODE and MTM1_OSE_FIT) were written in Python 3.16 (python.org) and are described as full code walkthroughs in the README files on the GitHub repository. The models are available on GitHub (https://github.com/LewPhan1/MTM1_ODE.git). Parameters used for the modelling are described in the table below.

| **Parameter** | **Value** | **Determination method** |
|---|---|---|
| Probe-PI(3)P Affinity | 0.457 µM | This paper **Figure S1 b** |
| Probe-PI Affinity | 50 µM | Assumed from ~100x lower binding then the specific interaction with PI(3)P |
| SNX17-PI(3)P Affinity | 3.16 µM | This paper **Figure S1 c** |
| MTM1-PI(3)P Affinity | 13.5 µM | Derived from ODE fit<sup>B</sup> |
| SNX17-Cargo Affinity | 2.2 µM | Literature<sup>C</sup> |
| MTM1-SNX17 Affinity | 60 µM | Literature estimate supported by ODE fitting<sup>D</sup> |
| Effective catalytic rate of MTM1 | 0.0024482 s<sup>-1</sup> | Derived from ODE fit<sup>A</sup> |
| Effective catalytic rate of MTM1/SNX17 dimer | 0.0024482 s<sup>-1</sup> | Assumed same as effective catalytic rate of MTM1 |
| Diffusion effect | N/A µM s<sup>-1</sup> | Unique fitting (described below)<sup>D</sup> |

#### Effective catalytic rate of MTM1 ^A^

It was assumed that the kcat of MTM1 in the conversion of PI(3)P to PI was constant between 0 mM NaCl and 150 mM NaCl. This assumption holds that the observed difference activities between these two salt concentrations (Fig 2 b and c) is the result of more MTM1 being driven to the membrane in the absence of salt. As was experimentally confirmed using fluorescently labelled, recombinant MTM1 (Fig 2e). Using fluorescently labelled recombinant MTM1 we determined that ∼23% of MTM1 was membrane-bound at 50 nM MTM1 and 0 nM NaCl. The affinity of MTM1 against PI(3)P was scaled in the ODE to achieve 23% binding under equivalent concentrations. The MTM1_ODE_FIT script was then used to fit 25 nM and 12.5 nM MTM1 simulation trajectories to experimental data (Fig. 2c). These two fits produced almost identical rate values (0.0024492 s^-1^ and 0.0024472 s^-1^) which were averaged to give the final value of 0.0024482 ^s-^1.

#### MTM1-PI(3)P affinity ^B^

Having gained an estimated fit for the effective catalytic rate of MTM1 and using the same assumption of equivalent kcat between 0 mM and 150 mM NaCl conditions, the MTM1_ODE_FIT script was used to estimate a value for the MTM1:PI(3)P affinity. The effective catalytic rate was fixed at 0.0024482 s^-1^, and the MTM1:PI(3)P affinity was then fitted against experimental data under 150 mM NaCl conditions. This produced an affinity estimate of 16.6 μM for the 50 nM MTM1 simulation and 10.4 for the 25 mM MTM1 simulation. The chosen affinity estimate was thus taken as the mean of these two values.

#### Literature derivation of Snx17-Cargo affinity ^C^

The 3D Kd for the affinity between SNX17 and the LRP1 cargo used in the dual channel experiments was taken as 2.2 μM from (Martín-González, et, al 2024). This value was then translated into a 2D for the ODE model using the method described in (Wu, et al, 2012) given that Kd values are typically described in three dimensions, and that this interaction occurs on the membrane where it is largely constrained to 2 dimensions. Briefly, this method involved multiplying the 3D Kd by the interaction’s confinement length (the height from the membrane where interaction between the molecules in confined). The confinement length was estimated using the measure tool in PyMol.

#### Derivation of SNX17-MTM1 affinity ^D^

The affinities for NPxY-cargo to SNX17 vary between 0.6 µM for human papillomavirus type 16 (HPV16) capsid protein L2 and 125 µM for the distal site of β1 integrin (ITGB1) (Martín-González, et, al 2024). An initial Kd of 60 µM was chosen for the SNX17:MTM1 interaction as this value sits approximately in the centre of this range. To confirm whether this was a reasonable assumption, the ODE solved for the SNX17:MTM1 affinity against an experiment containing 300 nM SNX17 (Fig. 3e). The model fit reproduced the observed activity increase from 0 nM SNX17 to 300 nM SNX17 using a Kd of 73.6 µM. Given this value was close to the initial 60 µM assumption, and a desire to avoid overfitting given experimental variability, the 60 µM assumed affinity was considered appropriate. This 3D affinity was converted into a 2D-equivilent parameter for the ODE model using the method described above (Wu, et al, 2012).

#### Unique fitting of diffusion effects ^E^

In order to fit the parameters of our ODE model to our experimental data, we have to take a delay of mixing and diffusion in our experimental setup into account. This becomes especially important for fast kinetic observed at low salt conditions. To account for this, a diffusion compartment was built into the model where components first enter the simulation. These then diffuse into the broader solution compartment via a unidirectional first-order rate equation. Delays in activity rates due to MTM1 diffusion can be observed from the lag time of the kinetics. Therefore, when constrained by fast kinetic activity, unique equilibration rates were fitted for the reaction.

#### Parameters for including VPS34 activity into ODE model

| Parameter | Value | Determination method |
|---|---|---|
| Maximum effective first-order rate coefficient for PI->PI(3)P conversion | 0.001 s<sup>-1</sup> | Assumed value for qualitative exploration of VPS34-mediated PI phosphorylation |
| Minimum effective first-order rate coefficient for PI->PI(3)P conversion | 0.00005 s<sup>-1</sup> | Assumed at 5% maximal activity to represent residual VPS34 activity under maximal feedback inhibition. |
| PI(3)P feedback midpoint | 40 nM | PI(3)P concentration at which VPS34 activity is halfway between its maximal and minimum values. Set to 50% of the simulated starting PI(3)P concentration. |
| Hill coefficient of PI(3)P-dependent VPS34 feedback | 2.0 | Assumed value for providing moderate non-linearity in the feedback response. |

---

**Figure S1:**
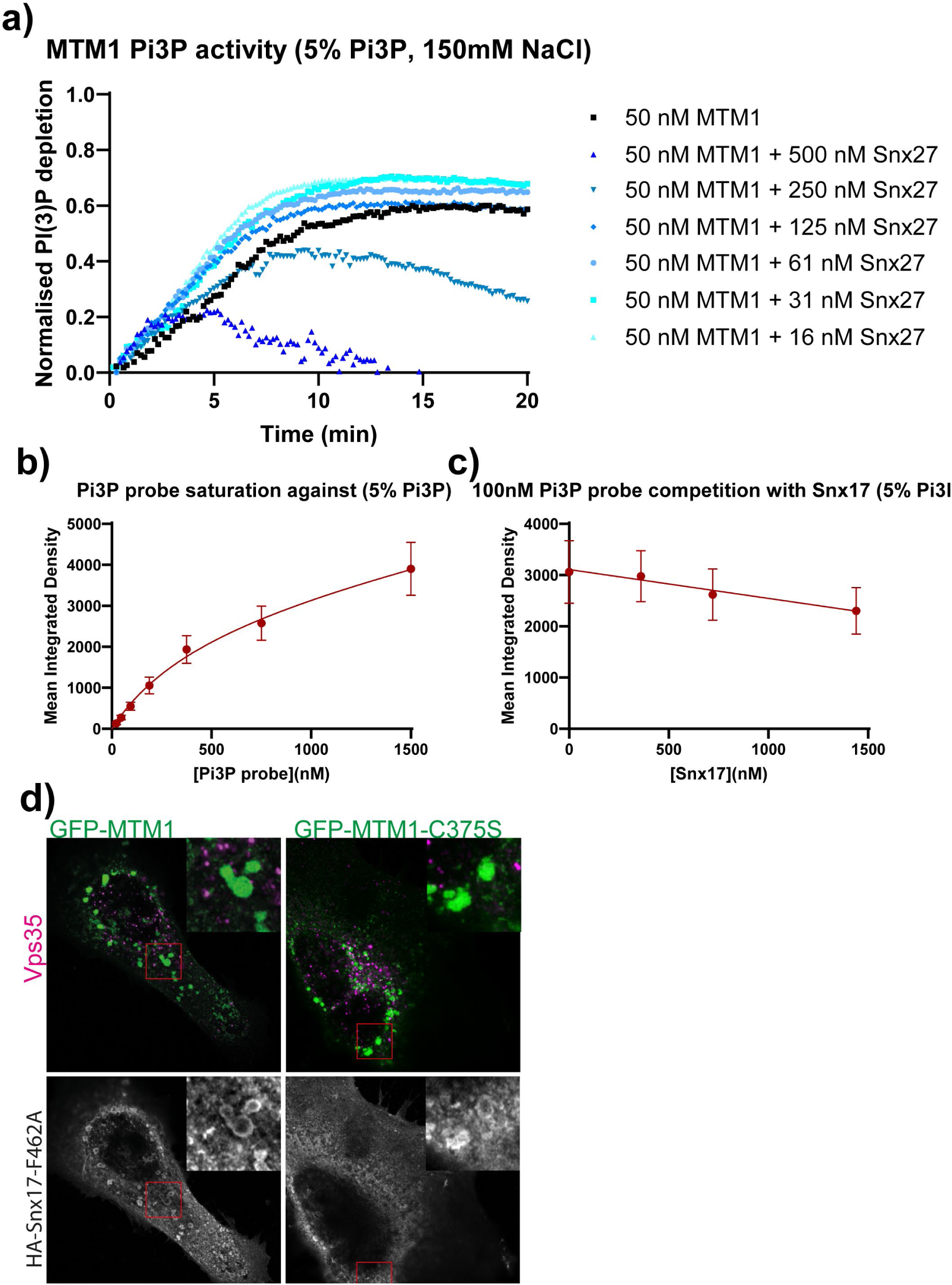
**a)** Phosphoinositide conversion assay was carried out in the presence or absence of a dilution series of recombinant SNX27 to measure activity of 50 nM MTM1. **b)** Affinity of dilution series of recombinant PI(3)P biosensor against membrane coated beads containing 5% PI(3)P. **c)** Competition binding assay of indicated recombinant SNX17 against 100 nM recombinant PI(3)P biosensor. **d)** HeLa cells were co transfected with either GFP-MTM1 or GFP-MTM1-C375S together with HA tagged SNX17-F462A

**Figure S2:**
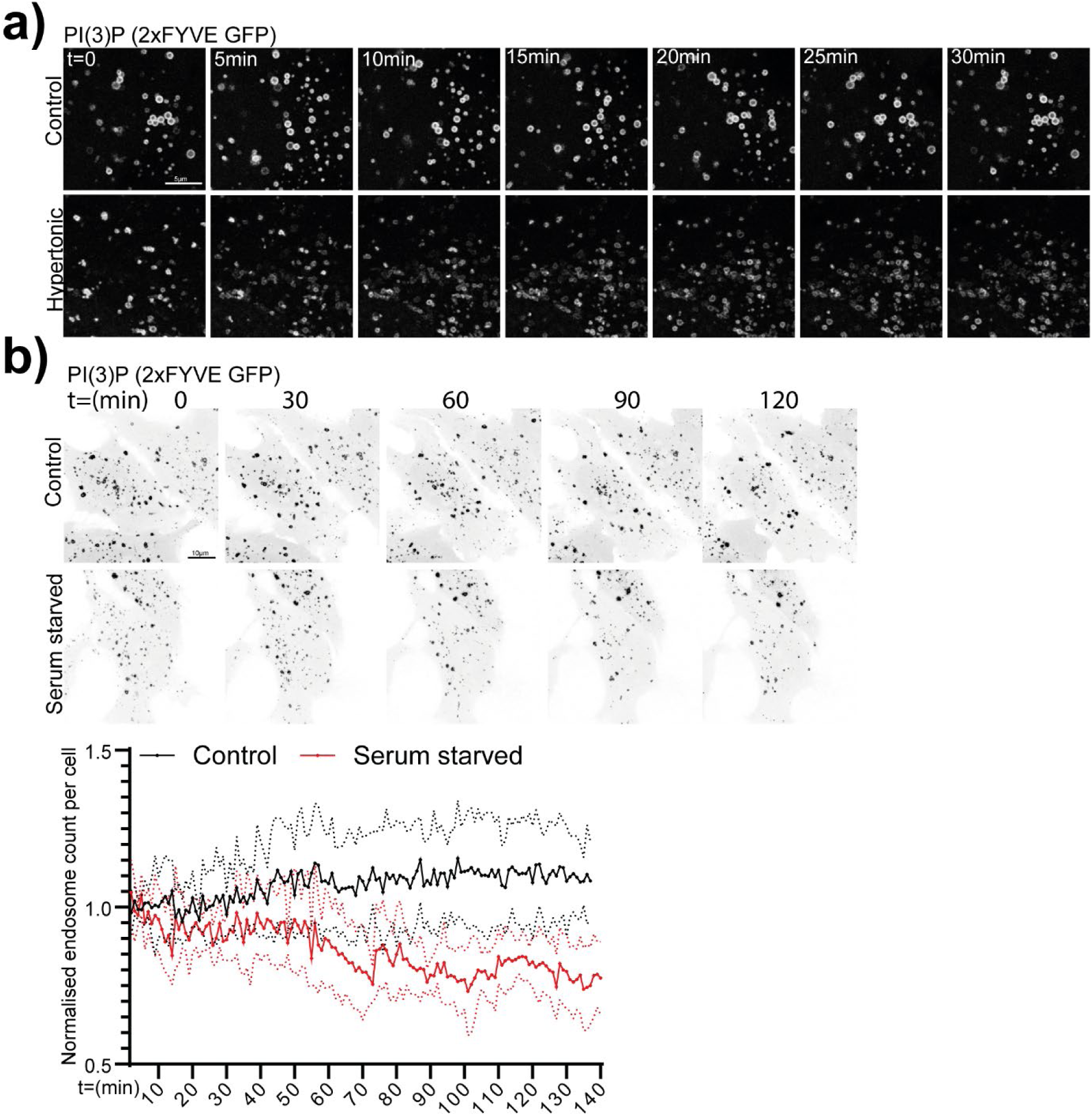
**a)** HeLa cells were transfected with 2xFYVE-GFP, treated with control or hypertonic media and imaged life for 30 minutes. **b)** HeLa cells expressing the 2xFYVE-GFP PI(3)P probe were imaged live at 37°C in CO2 independent media either in the presence or absence of Serum for 140 minutes at 1 complete Z volume every minute. Endosomes were segmented and normalised count from 6 individual cells was plotted over time ±SD

